# Stenoparib, an inhibitor of cellular poly (ADP-ribose) polymerase (PARP), blocks replication of the SARS-CoV-2 human coronavirus *in vitro*

**DOI:** 10.1101/2020.11.12.380394

**Authors:** Nathan E. Stone, Sierra A. Jaramillo, Ashley N. Jones, Adam J. Vazquez, Madison Martz, Lora M. Versluis, Marlee O. Raniere, Haley E. Nunnally, Katherine E. Zarn, Roxanne Nottingham, Jason W. Sahl, David M. Wagner, Steen Knudsen, Erik W. Settles, Paul S. Keim, Christopher T. French

## Abstract

By late 2020, the coronavirus disease (COVID-19) pandemic, caused by SARS-CoV-2 has caused tens of millions of infections and over 1 million deaths worldwide. A protective vaccine and more effective therapeutics are urgently needed. We evaluated a new PARP inhibitor, stenoparib, which was recently advanced to Stage II clinical trials for treatment of ovarian cancer, for activity against human respiratory coronaviruses, including SARS-CoV-2, *in vitro*. Stenoparib exhibits dose-dependent suppression of SARS-CoV-2 multiplication and spread in Vero E6 monkey kidney and Calu-3 human lung adenocarcinoma cells. Stenoparib was also strongly inhibitory to the HCoV-NL63 human seasonal respiratory coronavirus. Compared to remdesivir, which inhibits viral replication downstream of cell entry, stenoparib impedes entry and post-entry processes as determined by time-of-addition (TOA) experiments. Moreover, a 10 μM dosage of stenoparib – below the approximated 25.5 μM half-maximally effective concentration (EC_50_), combined with 0.5 μM remdesivir suppressed coronavirus growth by more than 90%, indicating a potentially synergistic effect for this drug combination. Stenoparib as a standalone or as part of combinatorial therapy with remdesivir should be a valuable addition to the arsenal against COVID-19.

**Importance:** New therapeutics are urgently needed in the fight against COVID-19. Repurposing drugs that are either already approved for human use or are in advanced stages of the approval process can facilitate more rapid advances toward this goal. The PARP inhibitor stenoparib may be such a drug, as it is currently in Stage II clinical trials for the treatment of ovarian cancer and its safety and dosage in humans has already been established. Our results indicate that stenoparib possesses strong antiviral activity against SARS-CoV-2 and other coronaviruses *in vitro.* This activity appears to be based on multiple modes of action, where both pre-entry and post-entry viral replication processes are impeded. This may provide a therapeutic advantage over many current options that have a narrower target range. Moreover, our results suggest that stenoparib and remdesivir in combination may be especially potent against coronavirus infection.

## Introduction

The novel SARS-CoV-2 coronavirus emerged from Wuhan, China in late 2019, and rapidly spanned the globe in a devastating pandemic (1). The disease, COVID-19, compromises the upper and lower respiratory systems and may affect all people (2). Although in many cases COVID-19 symptoms may be mild, some patients present with pulmonary distress leading to severe lung damage, and treatment options are limited (1, 3–5). Mortality estimates range from 0.5% to more than 5% (6). According to the Johns Hopkins COVID Resource Center (7), as of Nov. 10, 2020, over 10 million infections and more than 240,000 deaths have occurred due to COVID-19 in the USA alone, and the pandemic continues (8). A protective vaccine may become available (1, 2), but unless sufficient immunity can be achieved in the population, COVID-19 has the potential to cause morbidity and mortality for years to come. Without a protective vaccine, COVID-19 has largely been controlled through non-pharmaceutical measures such as quarantine, social isolation, and the use of personal protective equipment. Clearly, more efficacious treatments are needed.

Individuals who contract COVID-19 are most commonly infected by person-to-person transmission, where inhaled droplets containing infectious virions are seeded into the respiratory tract (1). The virions bind to respiratory epithelium via the affinity of the virus spike (S) complex to the angiotensin converting enzyme 2 (ACE2) receptor (9). A cellular serine protease TMPRSS2 plays a pivotal role in S protein priming (10), which in turn facilitates fusion between the viral and cellular plasma membranes and internalization of the virus-receptor complex by endocytosis. Subsequently, the virus is uncoated and releases its single-stranded RNA genome, which is processed, translated, and replicated in the host cytosol. Copies of the viral genome are packaged into bilayer membrane envelopes, and these new infectious virions are exported from the cell (11, 12). The SARS-CoV-2 lifecycle is typical of other coronaviruses, including the highly virulent SARS-CoV, the cause of Severe Acquired Respiratory Syndrome (SARS) (9). Conservation of key steps in the coronavirus viral lifecycles potentially constitutes an ‘Achilles Heel’ that is broadly susceptible to therapeutic intervention.

Antiviral therapeutics impede interactions between the virus and the host cell. Potential targets include the virus binding to the cellular receptor, viral entry or virus-host membrane fusion, viral transcription, translation, replication and export. (1). Stenoparib is an investigational, orally-available small molecule that inhibits poly (ADP-ribose) polymerase (PARP), a key enzyme in DNA repair (13). Stenoparib is unique in that it has dual inhibitory activity against the PARP 1/2 and tankyrase 1/2 enzymes, which are important regulators of the canonical Wnt/β-catenin checkpoint that is often dysregulated in metastatic breast cancer (14). Until August 2020, stenoparib was known as “2X-121” and previously as “E7449". Recently, another PARP inhibitor mefuparib (CVL218) was shown to inhibit SARS-CoV-2 *in vitro*. CVL218 suppressed SARS-CoV-2 infection in Vero E6 African Green Monkey cells (15). As implied by molecular modeling studies, CVL218 and other PARP inhibitors may block viral replication by interfering the viral nucleocapsid (N) protein binding to an RNA template (15).

The practice of repurposing existing drugs for new indications has advantages over developing an entirely new drug (16, 17). There are numerous repurposed drugs in use, including zidovudine, which was repurposed from the treatment of cancer to treat HIV/AIDS; the epilepsy drug topiramate, which is used to treat obesity; aspirin for analgesia and the prevention of colorectal cancer, among other examples (16). With repurposing, the risk of failure is lower compared to developing a new drug, because safety trials have already been completed and the *in vivo* pharmacokinetics have been characterized; thus, cost and time of development are reduced.

Moreover, the repurposing endeavor itself may reveal new disease targets and pathways. Altogether, reproposing can produce more rapid and efficient returns (16, 18). Stenoparib is currently in Phase II clinical trials for the treatment of ovarian cancer (14). Based on the recent promising results of the PARP inhibitor CVL218 against SARS-CoV-2 *in vitro* (15), we evaluated the activity of stenoparib against SARS-CoV-2, with an eye toward its use as a treatment for COVID-19.

## Results

### Stenoparib inhibits the replication of SARS-CoV-2 in Vero E6 cells

Based on the reported antiviral activity of other PARP inhibitors on SARS-CoV-2 *in vitro* (15) we aimed to determine whether stenoparib possessed a similar activity against the virus. Stenoparib was prepared as a solution in dimethylsulfoxide (DMSO) and used to treat Vero E6 cells infected with SARS-CoV-2 (USA-WA1/2020). Vero E6 is a common cell platform for propagating coronaviruses, including SARS-CoV-2 and SARS-CoV (19, 20). At 48 hours (h) after infection, RNA was isolated from cell supernatants, and viral copy number was estimated by reverse-transcription quantitative real-time polymerase chain reaction (RT-qPCR). Viral RNA measurements were compared to untreated cell controls, and to infected cells treated with a cocktail of camostat mesylate and E64d (‘C/E’), which are protease inhibitors that impede processing of the virus spike protein and interfere with its entry into the cell (10, 21). In parallel, we assessed potential toxic effects of stenoparib using the lactate dehydrogenase release assay, which indicates cytotoxicity due to cell lysis. As shown in **Fig. 1**, stenoparib demonstrated dose-dependent activity against SARS-CoV-2 at concentrations up to 30 μM with negligible cytotoxicity. The significant 63.2% reduction in viral load following treatment (t=8.608, P=0.0010) is comparable to the results reported for the CVL218 PARP inhibitor (35.2-99.7% inhibition) at comparable concentrations (15).

**Fig. 1.**
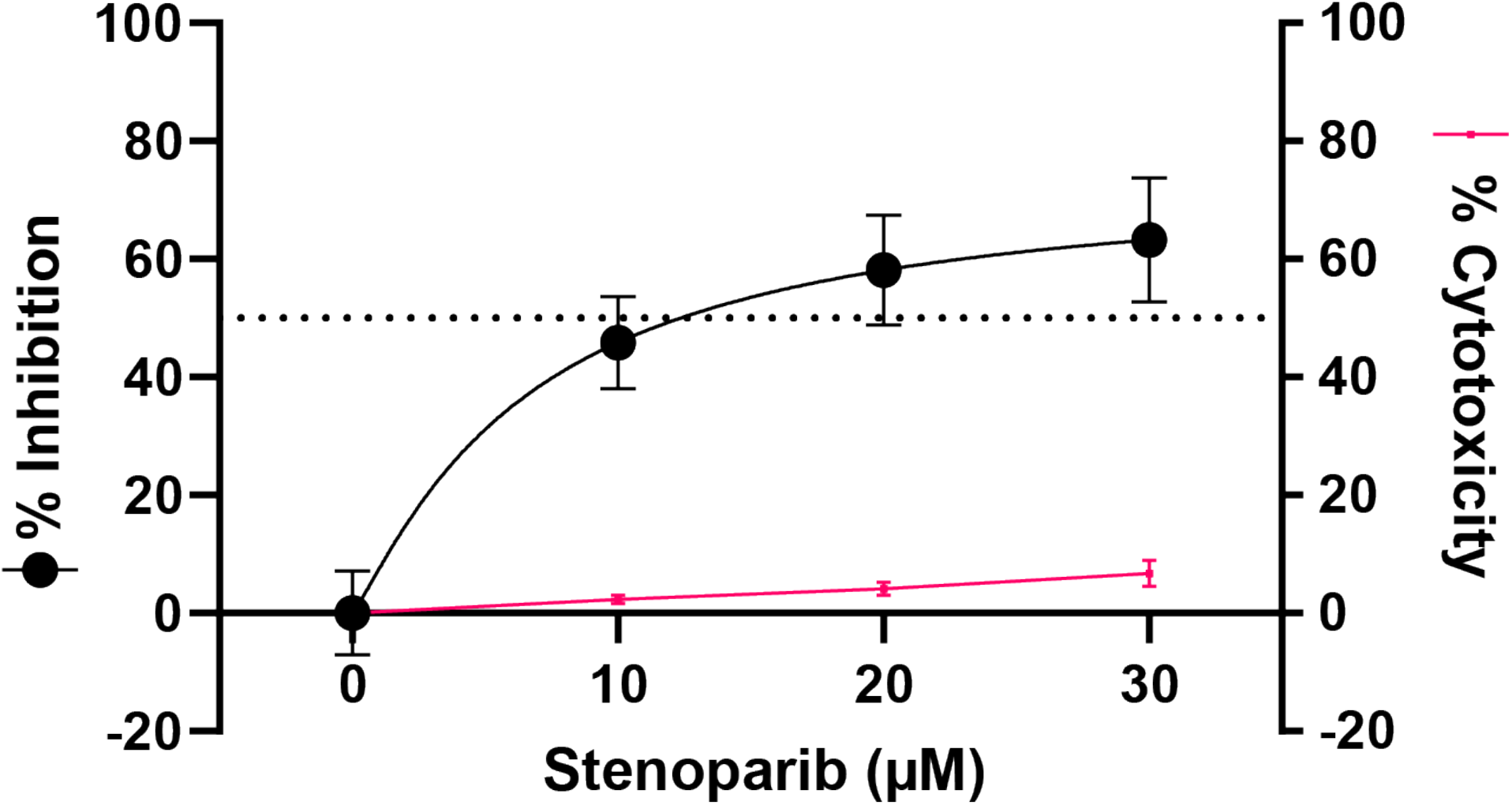
Stenoparib exhibits dose-dependent inhibition of SARS-CoV-2 as measured by RT-qPCR. RT-qPCR was performed on viral RNA collected from cell culture supernatants at 48 h post infection. Replicates within each run were averaged, and a total of three experiments were performed. Error bars were based on averaged standard deviations within runs. Cytotoxicity against Vero E6 cells was determined at 48 h using the Promega™ CytoTox 96™ assay kit, and values represent the average of the two independent experiments (reported in Fig. 2A).

At concentrations higher than 30 μM, and treatment durations exceeding 48 h, stenoparib displayed marked cytotoxicity to Vero E6 cells **(Fig. 2A),** which limited our capacity to comprehensively test the activity of the drug. We used a stenoparib response software predictor to pre-assess the susceptibility of human tumor cells based on the quantitative activity of 414 genes (22). When applied to human cell lines used with SARS-CoV-2 (23), the algorithm predicted that LLC-MK2 cells would be less sensitive to stenoparib toxicity than Vero E6 cells. The cell line Calu-3, originally isolated from the pleural effusion of a patient with lung adenocarcinoma (24), was predicted to be even more resistant than LLC-MK2. We verified this prediction for Calu-3 cells using stenoparib concentrations up to 60 μM, and exposure for up to 120 h, with no elevation in cytotoxicity over baseline conditions (t=8.237, P=0.0144) **(Fig. 2B)**.

**Fig. 2.**
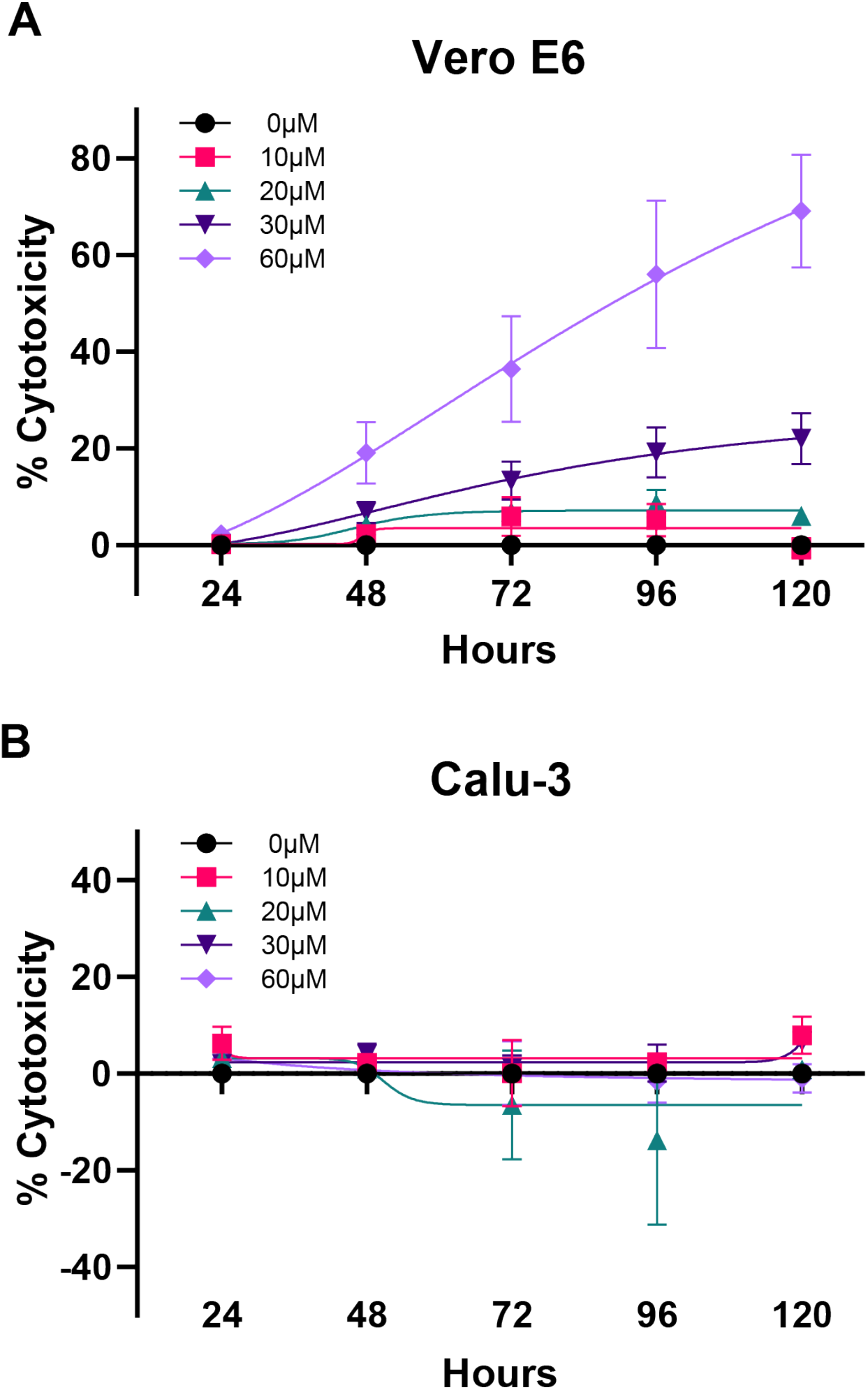
Stenoparib is cytotoxic in Vero E6 cells at concentrations greater than 30 μM, but not in Calu-3 cells. Cytotoxicity was determined using the Promega™ CytoTox 96™ lactate-dehydrogenase-release assay kit by harvesting culture media every 24 h up to 120 h post exposure. **A)** Vero E6 cells; **B)** Calu-3 cells, with 10, 20, 30, and 60 μM of stenoparib. Measurements were normalized to cells treated with 1.0% Triton X-100 and compared to untreated controls. Biological replicates from two runs were averaged, and median values are plotted. Results are representative of two experiments, and error bars are based on the standard deviation.

### Calu-3 lung epithelial cells as a platform for SARS-CoV-2

Since Calu-3 cells were more resistant to the toxic effects of stenoparib than Vero E6 cells, we utilized Calu-3 cells to test the effectiveness of higher doses than were previously achievable with Vero E6. The viral plaque assay comprehensively assesses inhibitors on the viral intracellular lifecycle, from virus entry, to multiplication, and cell to cell spread. Plaques result from cell damage and death following infection, appearing as empty regions or “dead zones” in the cell monolayer (25). Fresh media containing 2% FBS, with or without stenoparib or C/E control inhibitors, was applied to confluent monolayers of Calu-3 cells, which were then infected with SARS-CoV-2 for 1 h. At this time the infection media was removed, and the cells were overlaid with a semisolid matrix consisting of cell growth medium in 1.2% low melting temperature agarose, with or without stenoparib or control inhibitors. At 120 h post infection, cells were fixed with paraformaldehyde, stained with crystal violet, and the number of plaques were visually counted. As shown in **Fig. 3A** (see also **Fig. S1**), treatment with 30 μM stenoparib resulted in a 30.6% reduction of plaque forming units (PFUs) per well compared to infected, untreated cells (t=3.054, P=0.0379). Using a higher dose of 60 μM stenoparib, we observed near complete inhibition of plaque formation (94.0%; t=10.24, P=0.0005) with no significant cytotoxicity (t=0.446, P=0.6992), approaching the effect of the C/E control inhibitor (Fig. S1). These observations are mirrored by the results from RT-qPCR, which showed an 80.6% reduction of viral copy number with 60 μM stenoparib (P<0.0001). These observations affirm the prediction that Calu-3 cells are more resistant to the effects of stenoparib than Vero E6 cells and are suitable hosts for SARS-CoV-2 *in vitro*. It is interesting to speculate that Calu-3 cells may exhibit a degree of resistance to conditions that can be rapidly toxic in other, more rapidly dividing cell lines, which warrants further exploration. Indeed, the *in vitro* doubling time of Calu-3 cells (>60 h) was notably slower than either the Vero E6 (~24 h) or the LLC-MK2 (~36 h) cell lines.

**Fig. 3.**
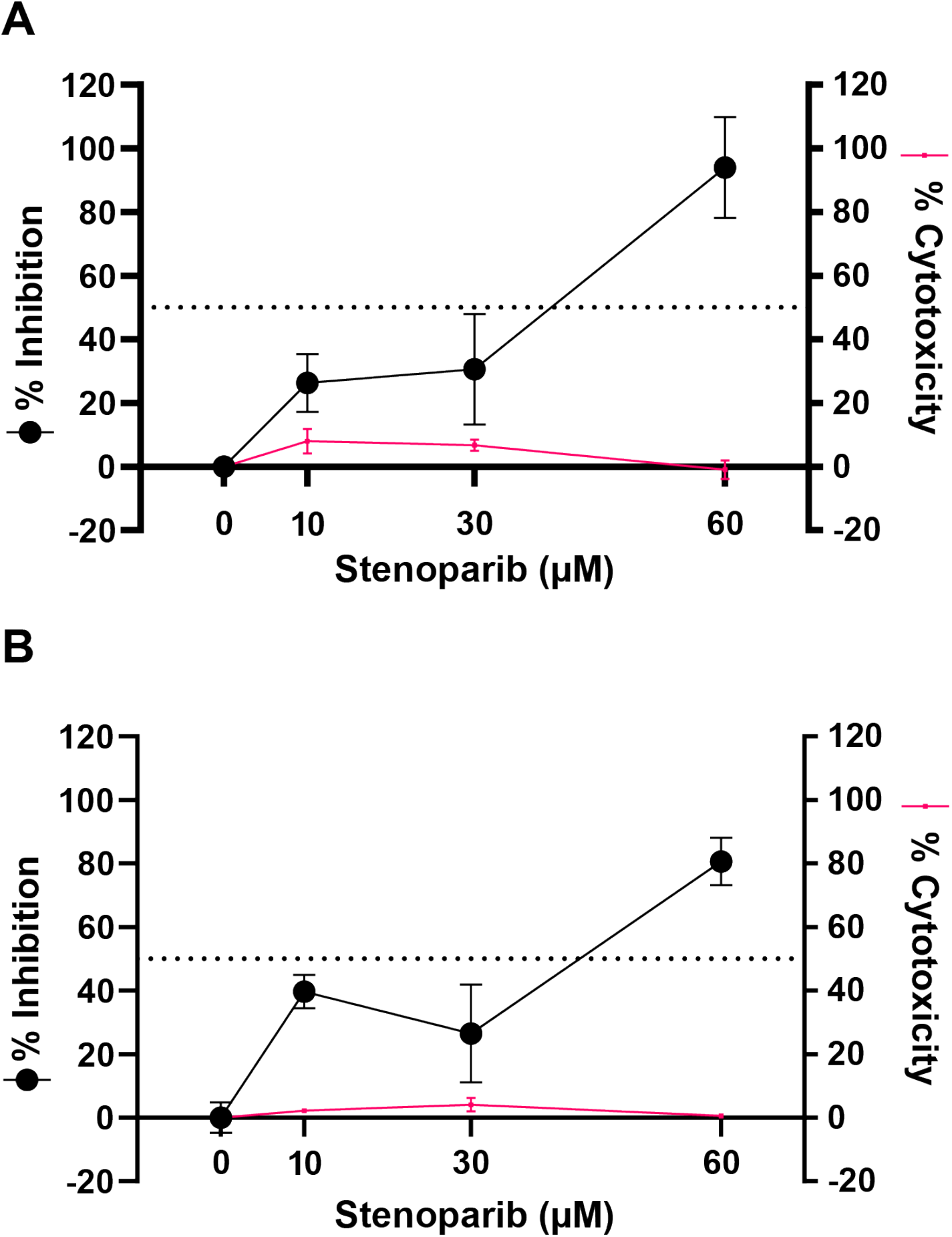
Stenoparib exhibits dose-dependent inhibition of SARS-CoV-2 in Calu-3 cells. **A)** Plaque forming efficiency using SARS-CoV-2. Values are normalized as a percentage of inhibition compared to infected, but untreated cells. Plaques were counted 120 h after infection, replicates from each run were averaged, and assays were performed three times. Error bars are based on the standard deviation across all runs. **B)** RT-qPCR was performed on viral RNA collected from cell culture supernatants at 48 h post infection and replicate values within each run were averaged; a total of three runs were performed. Error bars are based on averaged standard deviations within runs. Cytotoxicity against Calu-3 cells was determined at 48 and 120 h, as appropriate, using the Promega™ CytoTox 96™ assay kit, and values represent the average of the two independent experiments (reported in Fig. 2B).

### The NL63 virus as a surrogate *in vitro* model

In addition to SARS-CoV-2, several other human coronaviruses, including SARS-CoV, interact with human cells via the ACE2 receptor, and multiply intracellularly utilizing similar pathways (9). This group includes the respiratory coronavirus HCoV-NL63 (or just ‘NL63’), which is a cause of the seasonal colds in humans. While symptoms are generally mild, NL63 infections can be serious in infants and immunocompromised individuals (26–28). Based on its relatedness to SARS-CoV-2, and to establish a surrogate system for use in Biosafety Level-2 (BSL-2) instead of BSL-3 laboratory conditions, we evaluated NL63 for testing the effects of stenoparib.

The NL63 virus (NR-470) was propagated in LLC-MK2 Rhesus macaque kidney cells (29). Viral replication levels were assessed by plaque assay and RT-qPCR as performed for SARS-CoV-2. Controls were infected/untreated cells, and infected cells treated with the C/E inhibitor cocktail. Overall, the effects of stenoparib on NL63 corroborated the results of our experiments with SARS-CoV-2. Treatment resulted in a dose-dependent decrease in virus replication, achieving a 69.3% and 95.8% reduction of plaquing efficiency and viral copy number with 30 μM stenoparib, measured by plaque assay and RT-qPCR, respectively, compared to untreated controls (t=7.982 and 12.82, P=0.0002 for both) **(Fig. 4A and Fig. 4B)**.

**Fig. 4.**
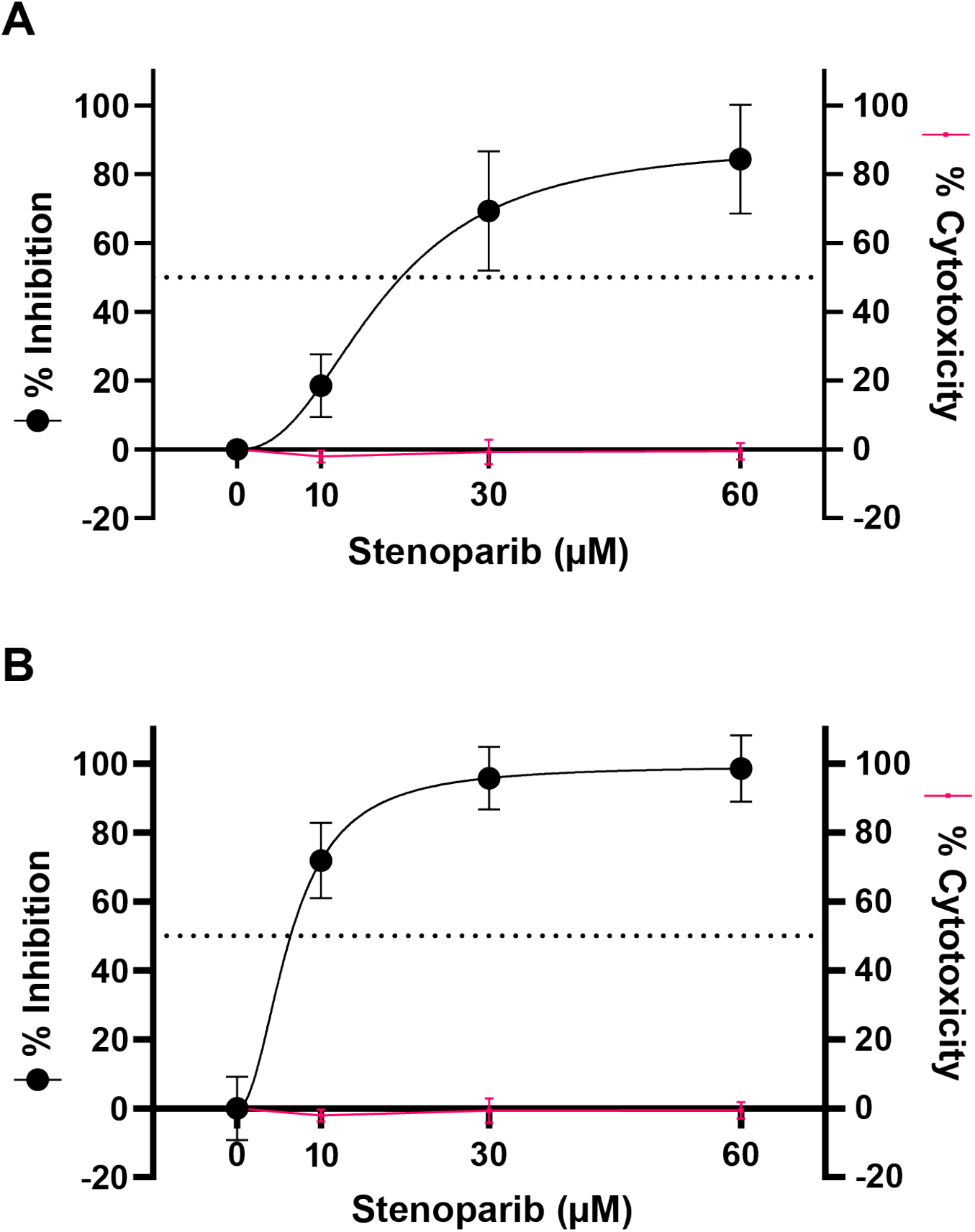
Stenoparib exhibits dose-dependent inhibition of HCoV-NL63 in LLC-MK2 cells. **A)** Plaquing efficiency values are normalized as a percentage of inhibition compared to infected, but untreated cells. Plaques were counted 120 h after infection, and assays were performed three times. Error bars are based on the standard deviation across all runs. **B)** RT-qPCR was performed on viral RNA collected from cell culture media at 120 h post infection. Biological replicates from each run were averaged, and three independent runs were performed. Error bars were based on averaged standard deviations within runs. Cytotoxicity against LLC-MK2 cells was determined at 120 h using the Promega™ CytoTox 96™ assay kit, and values represent the average of the three independent experiments.

### Identifying effects of stenoparib on the coronavirus lifecycle

Coronavirus inhibitors may target one or several intracellular growth stages, including virus entry (camostat mesylate, hydroxychloroquine), endosomal processing (hydroxychloroquine, rapamycin), translation and RNA processing (lopinavir), and transcription and replication (remdesivir) (1, 11, 30). By altering the time of addition (TOA) and duration of treatment *in vitro*, we can discern whether a drug affects virus entry, intracellular growth, or both. TOA experiments were conducted using the viral plaque assay with NL63 as a surrogate for SARS-CoV-2. RT-qPCR was performed in parallel to measure viral loads. We used remdesivir as a reference inhibitor, since its mechanism, target and dosage range are known (31). Experiments to determine the lifecycle stages affected by stenoparib were performed as follows: **a)** to ascertain the effect on virus entry, cells were transiently exposed to compounds starting 1 h before infection, and ending 1 h after infection; **b)** for effects on post-entry events, including transcription, processing, translation or replication, compounds were added 1 h after infection, when a number of virions would have already entered cells, and treatment was continued until the experimental endpoint at 120 h; **c)** to examine the maximum achievable effect of the compounds, a full-time assay was performed. Treatment was initiated simultaneously with virus infection and continued until the 120 h endpoint (see **Methods** for a detailed description).

As shown in **Fig. 5A,** the antiviral activity of stenoparib is most notable when added post-infection. The 60 μM stenoparib dose achieved complete inhibition of NL63 plaquing, with no detectable (N/D) plaques following post-entry treatment compared to untreated cells (P<0.0001). This is on par with the effect of 4 μM remdesivir, which also eliminated plaque formation. Likewise, assay results for the full-time treatment were comparable between stenoparib and remdesivir, with 88.4% reduction in plaque efficiency for stenoparib (t=5.582, P=0.0051), and full inhibition for remdesivir vs. untreated cells. This is consistent with our results from RT-qPCR, where stenoparib produced 98.7% (t=9.988, P=0.0099) and 95.5% (t=9.663, P=0.0105) inhibition vs. untreated controls for post-entry and full-time drug exposure, and also comparable to the activity of remdesivir. These data are in line with those reported previously for the CVL218 PARP inhibitor and remdesivir (15).

**Fig. 5.**
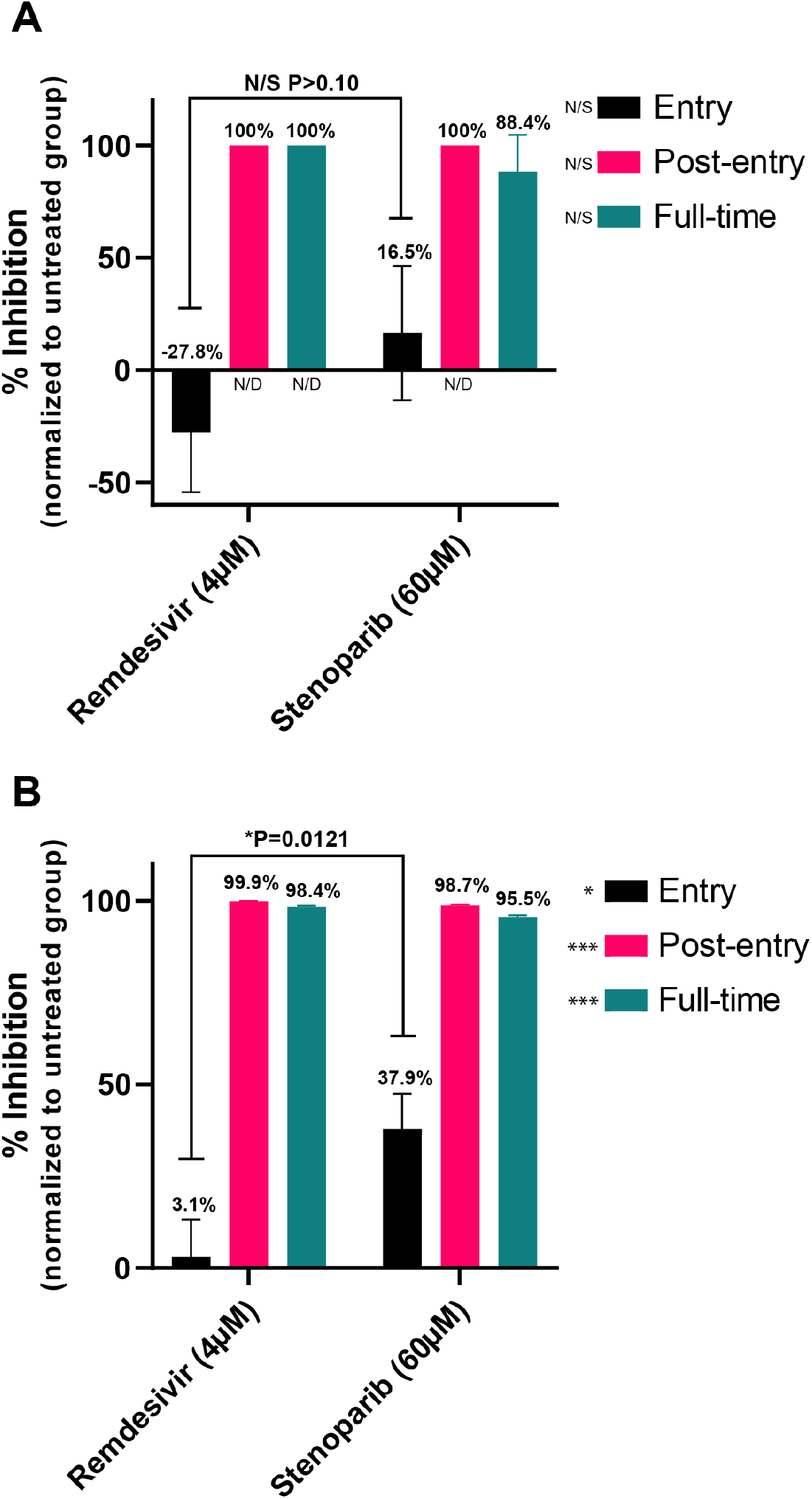
Stenoparib inhibits HCoV-NL63 entry and post entry events while remdesivir inhibits post-entry. **A)** Plaque assays were performed three times and replicate plaque forming unit counts from each run were averaged. Error bars are based on standard deviation among runs. Brackets indicate the t-test comparison and P-value for the “Entry” group. No significant difference was observed between stenoparib and remdesivir under any treatment. N/D indicates that no plaques were detected in these treatment groups. **B)** RT-qPCR was performed on viral RNA collected from cell culture media at 120 h post infection and replicate values within each run were averaged; a total of three runs were performed. Error bars were based on averaged standard deviations within runs. Brackets indicate the t-test comparison and P-value for the “Entry” group. Significant differences were observed between stenoparib and remdesivir under all treatments.

With the plaque assay, we noticed a 16.5% reduction in plaque formation following transient treatment with stenoparib early in the infection time course (i.e. ‘Entry’; **Fig. 5A**). This effect was not markedly different than the results recorded for remdesivir (−27.8%; t=1.919, P=0.1275). Inhibition of virus entry is not expected for remdesivir since its activity is specific to blocking of RNA replication (31, 32), which is a mid-late event in the viral lifecycle. In contrast, our results from RT-qPCR strongly support a specific effect for stenoparib on inhibiting virus entry, where a 37.9% reduction in viral load is observed in the entry assay compared to just a 3.1% reduction for remdesivir (t=4.352, P=0.0121) (**Fig. 5B**). Effects on viral entry are consistent with the predicted activity of stenoparib on processes involved in early coronavirus infection events (**see Discussion**). Taken as a whole, these observations suggest that stenoparib may affect multiple targets that play roles in the early and late stages of coronavirus intracellular multiplication.

### The combination of stenoparib and remdesivir strongly inhibits NL63

Combination drug therapies are widely used for the treatment for some of the worst human diseases, including cancer (33), HIV/AIDS (34), and multidrug-resistant tuberculosis (35). The strategy of combination therapy seeks to increase the beneficial effects of multiple drugs, lower their doses to reduce adverse effects, and minimize the induction of resistance (36). Generally, the activity of a drug combination is considered additive when the combined effect of two drugs is equivalent to their individual doses, while if the effect is less than additive, the combination is considered antagonistic. Synergy occurs when the combined effect is greater than the additive effect (36). Combinations of two or more drugs may lead to a synergistic effect by combining different mechanisms of action (MOA). Examples of synergistic combinations of drugs with distinct MOAs include streptomycin–penicillin (37), trimethoprim–sulfa drugs (Co-trimoxazole) (38), and β-lactam/β-lactamase inhibitor combinations such as amoxicillin-clavulanate (39) against bacterial infections.

Based on the fact that stenoparib and remdesivir inhibit coronavirus by distinct MOAs (31, 32, 40), we approximated the half maximal effective concentration (EC_50_) of stenoparib and remdesivir as 25.5 μM and 0.46 μM using the NL63 virus and plaque assays. Calculations were aided by the online calculator from AAT Bioquest (“Quest Graph™ EC_50_ Calculator.” *AAT Bioquest, Inc*, 26 Oct. 2020, https://www.aatbio.com/tools/ec50-calculator). We hypothesized that a combination of stenoparib and remdesivir would be more potent than the individual compounds. To test this, we combined a range of doses of stenoparib with 0.5 μM remdesivir and tested these for activity against NL63.

As shown in **Fig. 6**, complete inhibition of NL63 plaque formation was achieved with 60 μM stenoparib, in line with our earlier results with SARS-CoV-2 and Calu-3 cells **(Fig. 3).** Complete inhibition was also achieved with 1.0 μM remdesivir. However, a combination of 10 μM stenoparib and 0.5 μM remdesivir was more effective than either compound alone at these doses, achieving 90.7% inhibition for the combination, vs. 18.5% inhibition for 10 μM stenoparib, and 65.6% for 0.5 μM remdesivir, suggesting at least additive effects when the drugs are combined. Notably, the stenoparib dose (10 μM) used in the combination is far below the compound’s EC_50_ of 25.5 μ<u>M.</u> Altogether, these results support investigating the use of stenoparib and remdesivir as a combinatorial therapy for SARS-family coronavirus infections.

**Fig. 6.**
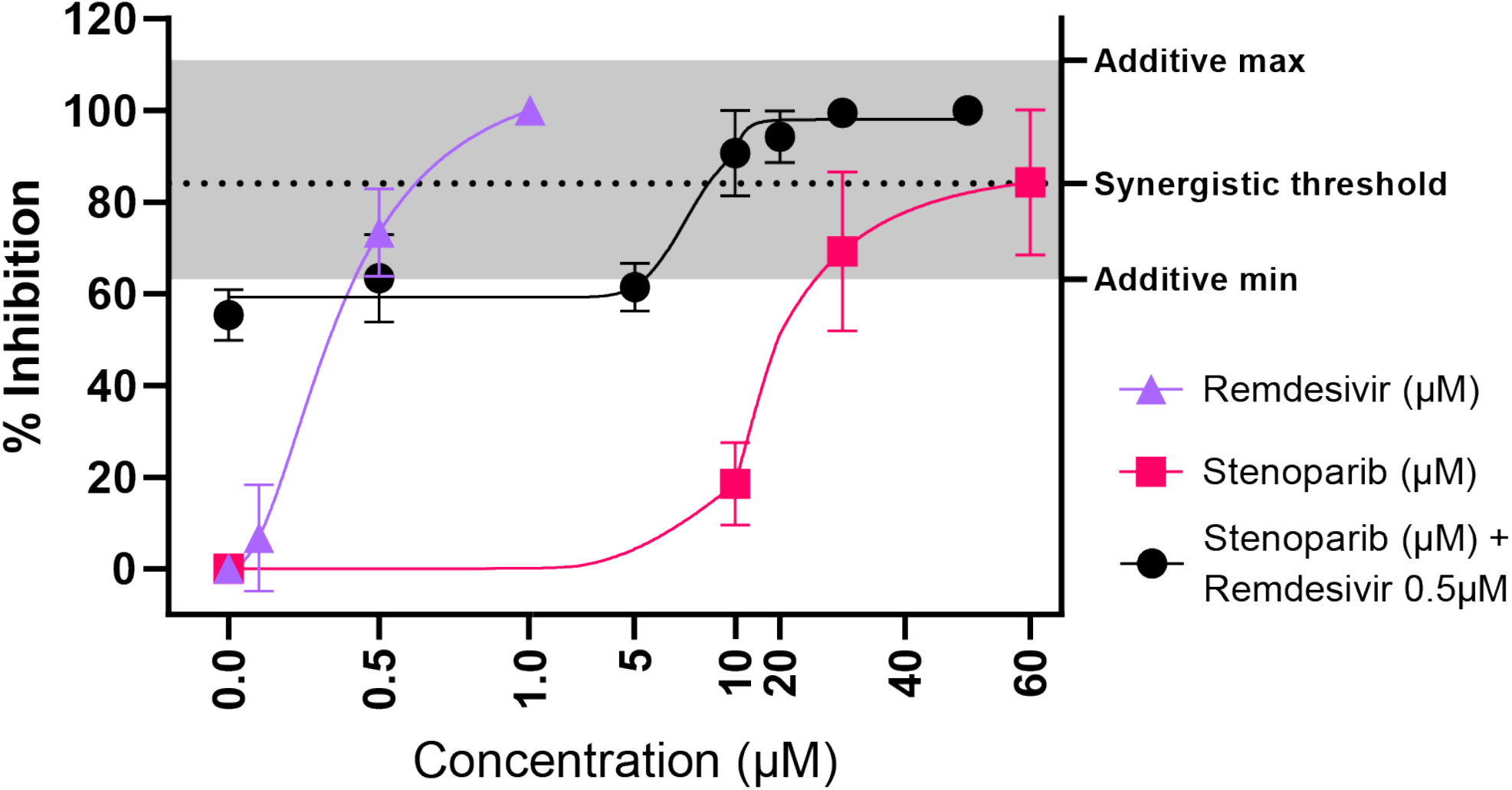
Stenoparib and remdesivir in combination is a potent inhibitor of NL63. Plaque assays were performed a minimum of two times and replicate values from each run were averaged. Plaquing efficiency values are normalized as a percentage of inhibition compared to infected, but untreated cells. Three data sets are plotted to illustrate the treatment of NL63 with 1) stenoparib and 2) remdesivir as monotherapies and as a 3) combination therapy, whereby increasing concentrations of stenoparib are combined with 0.5 μM remdesivir; the EC_50_ was computationally approximated at 0.46 μM. The stenoparib monotherapy data is the same as previously reported (Fig. 4A). The synergistic activity threshold is defined as the sum of the mean values of 10 μM stenoparib and 0.5 μM remdesivir as monotherapies, while the grey highlighted area represents the minimum and maximum possible additive activity values based on the range of error for those same concentrations observed during these experiments.

## Discussion

Prior to the emergence of COVID-19, attempts to identify inhibitors of coronavirus were mainly focused on SARS- and MERS-CoVs. The recent efforts to develop COVID-19 therapeutics spans the gamut from new drug discovery to repurposing existing drugs. There are some excellent reviews on the subject (17, 41). Here, we focus on the compound stenoparib, formerly known as 2X-121, an inhibitor of the cellular enzyme PARP-1/2. Stenoparib is thought to inhibit SARS-CoV-2 by multiple mechanisms, predominately by inhibiting of ADP-ribosylation of proteins required for virus replication and assembly (42). ADP-ribosylation is a conserved, post-translational modification that is key for proper formation of the coronavirus nucleocapsid, and inhibition can negatively affect packaging of the viral genome and virion stability. Specific targets of ADP-ribosylation include viral nsp3 protein, which is essential for virulence and a component of the replication/transcription complex (RTC) (43, 44). Moreover, PARP inhibitors may exert additional protective effects at the host and cellular levels by reducing depletion of NAD+ and ATP that leads to cell necrosis (40), as well as decreasing the pro-inflammatory NF-kB-triggered cytokine storm that can damage host organs (45). It has also been suggested that PARP inhibitors enhance the degradation of the host type I interferon receptor (IFN-1R), which would also have a modulatory effect on the host inflammatory response (46).

In this study, inhibition of SARS-family coronaviruses by stenoparib *in vitro* is likely due to interference with multiple stages of the virus lifecycle. Consistent with its predicted mechanism, stenoparib is effective when introduced post-infection. Additionally, we noted an association between stenoparib treatment and decreased virus counts soon after infection. This may reflect activity against multiple targets, including those involved in promoting virus entry. Stenoparib interference with the activity of tankyrase and Wnt/β-catenin signaling can downregulate expression of the ACE2 receptor, leading to a decrease in the number of viruses that can enter the cell, which is consistent with effects on virus entry and with our observations. In contrast, inhibition by remdesivir was predominately on post-entry events – its effect on virus entry was minimal at best, which is in line with its specific MOA of targeting the virus replication machinery. Overall, our observations imply multiple mechanisms for stenoparib, including impeding of viral entry and intracellular growth via modifications of multiple viral and host proteins.

Other human coronaviruses that utilize ACE2 for binding and entry may be suitable as surrogate platforms for the study of SARS-CoV-2 *in vitro,* so long as they can be propagated in the laboratory and are able to elicit cellular infection phenotypes that can be quantitatively measured. We show that the human seasonal coronavirus NL63, which can cause a cold-like illness in humans (26), is such an example. Like SARS-CoV-2 and SARS-CoV, NL63 is internalized following the binding of viral S-complex proteins to the ACE2 receptor (28, 47–49). According to our observations, NL63 is susceptible to inhibition by compounds that affect SARS-CoV-2, including remdesivir, the protease inhibitors camostat mesylate and E64d, and stenoparib, the subject of this study.

In light of the *in vitro* cytotoxic effects of stenoparib, we were unable to test it in Vero E6 cells against SARS-CoV-2 at doses exceeding 30 μM. Stenoparib was developed as a cytotoxic drug for cancer treatment (14), so it is not surprising that it showed dose-dependent cytotoxicity against rapidly-dividing Vero E6 cells. PARP inhibitors typically express their lethality after several replication cycles (50). The susceptibility of Vero E6 cells to stenoparib toxicity may be linked to its fast-growing phenotype, since two replication cycles can be achieved in as little as 48 h. Although LLC-MK2 cells, which are utilized for propagation and testing of NL63 are substantially more resistant to stenoparib, their susceptibility to SARS-CoV-2 infection is suboptimal. We were thus presented with a conundrum in how to test high doses of stenoparib against SARS-CoV-2 to assess its maximum activity. This was addressed through the use of the Calu-3 human lung carcinoma cell line to model SARS-CoV-2 infections. Our results confirm the suitability of Calu-3 as a host for SARS-CoV-2 and its high degree of resistance to stenoparib toxicity. This characteristic of Calu-3 cells was predicted *in silico* using a method previously employed on clinical tumor biopsies (22), and experimentally validated in this study. Infection of Calu-3 monolayers with SARS-CoV-2 formed large, clearly visible plaques, and in this regard, the performance of Calu-3 surpassed Vero E6. A notable characteristic of Calu-3 is its slow growth properties, which may coincide with a general degree of resistance to compounds that target essential cellular pathways and are toxic to more rapidly dividing cells.

A promising drug to emerge from COVID therapeutic trials is the nucleoside analog remdesivir, which shows activity against phylogenetically diverse viruses including Ebola, Nipah, respiratory syncytial virus (RSV), and coronaviruses such as SARS- and MERS-CoV (51), SARS-CoV-2 (31) and, as reported here, the seasonal human HCoV-NL63. Reviews on the MOA of remdesivir have been published (17, 41, 51). After entering the cell, remdesivir is triple phosphorylated to form remdesivir-triphosphate, and this structure is thought to inhibit RNA polymerase, resulting in chain termination. While the precise molecular mechanism is not clear (51), the EC_50_ of remdesivir has been reported to be 0.77 μM for SARS-CoV-2 (31), which is in line with our measurement of 0.54 μM for SARS-CoV-2, and comparable to our experimentally-determined EC_50_ of 0.46 μM against NL63.

While remdesivir inhibits the viral replicon, our data support multiple targets for stenoparib. Moreover, stenoparib and remdesivir may be a potent combination at inhibiting SARS-family coronaviruses inside cells. A mixture of 10 μM stenoparib and 0.5 μM remdesivir was more successful at inhibiting the NL63 virus than either compound alone at these doses. Considering their distinct mechanisms and high potency, a combination of remdesivir and stenoparib is likely to produce a synergistic effect on additional SARS-family coronaviruses, including SARS-CoV-2. Studies involving this combination in susceptible COVID-19 animal models are in line for efficacy testing.

## Materials and Methods

The antiviral activity of stenoparib *in vitro* was assessed against the novel coronavirus SARS-CoV-2 isolate USA-WA1/2020 (BEI Resources, NIAID, NIH: SARS-Related Coronavirus 2, Isolate USA-WA1/2020, NR-52281) and human coronavirus strain HCoV_NL63 (BEI Resources, NIAID, NIH NR-470). We used Vero E6 (ATCC CRL-1586) and Calu-3 cells (ATCC HTB-55) from the American Type Culture Collection (ATCC, Manassas, VA, USA) in EMEM (Eagle’s Minimum Essential Medium) supplemented with 2% or 10% fetal bovine serum (FBS), 100 U/mL penicillin, 100 μg/mL streptomycin ‘pen-strep’, 0.01 M HEPES, 1 mM sodium pyruvate, 1x non-essential amino acids solution (SH3023801, Thermo Fisher), and 2 mM L-glutamine, for the propagation and experimentation with SARS-CoV-2. LLC-MK2 cells (ATCC CCL-7), maintained in Medium 199 (M4530, Millipore Sigma) supplemented with FBS and pen-strep, were used for the experiments with the NL63 coronavirus. Inhibition of viral replication was assessed using reverse-transcription quantitative real-time PCR (RT-qPCR) to measure the number of virions released into the cellular supernatant.

### Plaque assays

6-well plates (CLS3516, Millipore Sigma) were seeded with ~3.0×10^5 cells/well and incubated for 48-72 h at 37°C in 5% CO_2_ until 80-90% confluency was reached. Calu-3 cells were incubated for >10 days to achieve 80-90% confluency. Prior to infection the media was replaced with fresh media containing 2.0% FBS with varying concentrations of stenoparib as appropriate for each experiment (see results) and infected with coronavirus (SARS-CoV-2 or NL63) at an MOI of 0.013 for SARs-CoV-2 and 0.003 for NL63. Media was then replaced with a 1 X Dulbecco’s MEM (DMEM - Millipore Sigma) / 1.2% low-melting agarose (Bio-Rad) overlay containing the appropriate drug concentration for each experiment. This was allowed to solidify at room temperature for 15 minutes, and incubated for 120 h at 37°C in a 5% CO2 atmosphere. A cocktail of the protease inhibitors camostat mesylate and E64d (C/E) was a control for all experiments (21). SARS-CoV-2 manipulations were conducted in a BSL-3 facility. 2.0 ml of 4% paraformaldehyde was added to each overlay for 30 minutes, followed by staining with 1.0% crystal violet, removal of the overlay, and a triple rinse with PBS. Plaque forming units (pfu) were counted, averaged, and normalized to the untreated control group. Each run contained three biological replicates and was conducted a minimum of two times. Standard deviation was calculated using the variation of averaged counts among all runs. Values were plotted using GraphPad Prism version 8.0.0 for Windows (GraphPad Software) and annotated using Adobe Illustrator (Adobe Systems Incorporated). Statistical significance was determined using a parametric unpaired t-test in GraphPad Prism version 8.0.0.

### RT-qPCR: Infection and viral RNA extraction

12-well plates (CLS3513, Millipore Sigma) were seeded with ~1.0×10^5 cells/well and incubated until 80-90% confluency was reached. Growth media was replaced and infected with coronavirus (SARS-CoV-2 or NL63) at an MOI of 0.04 for SARs-CoV-2 and 0.01 for NL63 for up to 120 h at 37°C in 5% CO2 atmosphere. C/E was used as a control inhibitor for all experiments. Each run contained two biological replicates and was conducted three times. 400 μL of supernatant was harvested at 48 h for SARS-CoV-2, 120 h for NL63. RNA was extracted using Invitrogen Pure-Link RNA kits (Thermo Fisher) according to their recommendations.

### RT-qPCR: Signature identification and qPCR assay design

Two TaqMan qPCR assays were designed for the SARS-CoV-2 or the NL63 coronaviruses. For NL63, the reference genome (NC_005831) was divided into 200 nucleotide fragments, which were aligned against a set of 2,771 coronavirus genomes with BLAT v36.2 (52) in conjunction with LS-BSR v1.2.2 (53). Regions were identified that had a BLAST score ratio (BSR) (54) value ≥0.8 in 60 NL63 genomes, and a BSR value <0.4 in all other coronavirus genomes. A total of 10 fragments were highly specific to all NL63 genomes. Primers and probes were identified using Primer3 v2.3.6 (55). A similar process was conducted for SARS-CoV-2 assay design, except the GCF_009858895.2 reference genome was used. A total of 4 fragments were unique to 64 distinct SARS-CoV-2 genomes. A probe was designed targeting the spike (S) protein furin cleavage site with Primer3.

The SARS-CoV-2 qPCR amplified a 125 bp region of the S protein using forward primer CoV2-S_19F (5’-GCTGAACATGTCAACAACTC-3’) and reverse primer CoV2-S_143R (5’-GCAATGATGGATTGACTAGC-3’) with MGB TaqMan probe CoV2-S_93FP (5’-ACTAATTCTCCTCGGCGGGC-3’) labeled with FAM dye, which was designed based off the SARS-CoV-2 genome GCF_009858895.2 (GenBank: MN908947.3), while the NL63 qPCR amplified a 191 bp region of a membrane protein (GenBank: YP_003770.1) using forward primer NL63_10F (5’-TGGTCGCTCTGTTAATGAAA-3’) and reverse primer NL63_200R (5’-AAATTTCTTCCTAGCAGCTC-3’) with MGB TaqMan probe NL63_102RP (5’-CCCTCCTGAGAGGCAACACC-3’), labeled with a VIC dye, which was based off the HCoV_NL63 genome (GenBank: MN306040.1).

### RT-qPCR: Reverse transcription and PCR amplification

We initially used a two-step method where viral RNA was converted into cDNA using Invitrogen SuperScript IV VILO Master Mix (11766500, Thermo Fisher) in 96-well format (18021-014, Thermo Fisher); in a SimpliAmp® thermocycler (Applied Biosystems). 1 μL of template cDNA was then subjected to qPCR in 10 μL reactions containing 1x TaqMan Universal Master Mix II (w/o AmpERASE UNG), with 0.2 μM of each forward and reverse primer and 0.1 μM of probe for the SARS-CoV-2 qPCR and 0.25 μM of each forward and reverse primer and 0.125 μM of probe for the NL63 qPCR). Amplification was performed in triplicate using either a QuantStudio 7 flex or QuantStudio 12K (Applied Biosystems) utilizing: 10’ at 95°C, then 40 X at 95°C 15”, 60°C for 1’. Another approach employed a one-step procedure in which viral RNA was converted to cDNA using the TaqMan primers, followed by qPCR (4x Reliance One-Step Multiplex RT-qPCR Supermix) with the same primers, probes, and concentrations as for the two-step approach. Triplicate reactions were performed using QuantStudio under the following conditions: 50°C for 10’; 95°C, 10’; 40 cycles of 95°C 10” and 60°C 30”. Positive amplification and non-template controls were included on every run.

### RT-qPCR: Data analysis

Synthetic double-stranded DNA fragments were generated (gBlocks Gene Fragments, Integrated DNA Technologies) as qPCR controls, and contained amplification primers for either SARS-CoV-2 or NL63 targets, and were elongated to 200 bp. These gBlocks were resuspended according to the manufacturer’s protocol and quantified using a Qubit 4 fluorometer with a dsDNA HS assay kit (Q32851 Thermo Fisher), then normalized to 10^8 copies per μL. Using serial dilution, we were able to extrapolate viral copy number in each of the experimental samples. Based on these standards, the QuantStudio instrument software generated a curve to quantify sample reactions. The calculated quantities for each sample were averaged, and the standard deviation was calculated among reactions. Values for the experimental replicates and the standard deviations among experimental runs were averaged, and then normalized to the untreated control group to obtain percent inhibition values. These were plotted using GraphPad Prism version 8.0.0 for Windows and annotations were added using Adobe Illustrator. Where appropriate, statistical significance was determined using a parametric unpaired t-test in GraphPad Prism version 8.0.0.

### Cytotoxicity

Cytotoxicity was measured using the Promega™ CytoTox 96™ Nonradioactive Cytotoxicity Assay kit (G1780) in 50 μL reactions according to the manufacturer’s protocol. The absorbance at 490 nm (A490) was measured using a BioTeK Synergy™ HT plate reader model # 7091000. Relative cytotoxicity was calculated by dividing the experimental LDH release as measured at 490nm by the maximum LDH release control multiplied by 100.

### Time of addition experiments

For the full-time experiments, virus, drug and cells were incubated for 1 h. Media was then replaced with fresh media containing the drug. Entry experiments were pretreated with drug for 1 h and then infected with virus for an additional hour, followed by media replacement that lacked drug. Post-entry experiments utilized cells that were infected with virus for 1 h, and media was replaced with fresh media containing the drug. Statistical significance was determined using a parametric unpaired t-test in GraphPad Prism version 8.0.0.

### Stenoparib in combination with remdesivir

We performed plaque assays, and used these data to estimate the EC_50_ of stenoparib and remdesivir (329511, MedKoo Biosciences, Morrisville, NC, USA) against NL63 (both drugs) and SARS-CoV-2 (remdesivir only) in LLC-MK2 and Vero E6 cells according to results from at least two experimental runs. The EC_50_ values were approximated with the aid of the online calculator from AAT Bioquest (“Quest Graph™ EC_50_ Calculator.” *AAT Bioquest, Inc*, 26 Oct. 2020, https://www.aatbio.com/tools/ec50-calculator).

## Acknowledgments

This research was supported by a grant from The Flinn Foundation and continuing support of The Cowden Endowment for Microbiology. Core experiments were supported by service fees paid by Allarity Therapeutics.

---

**Fig. S1.**
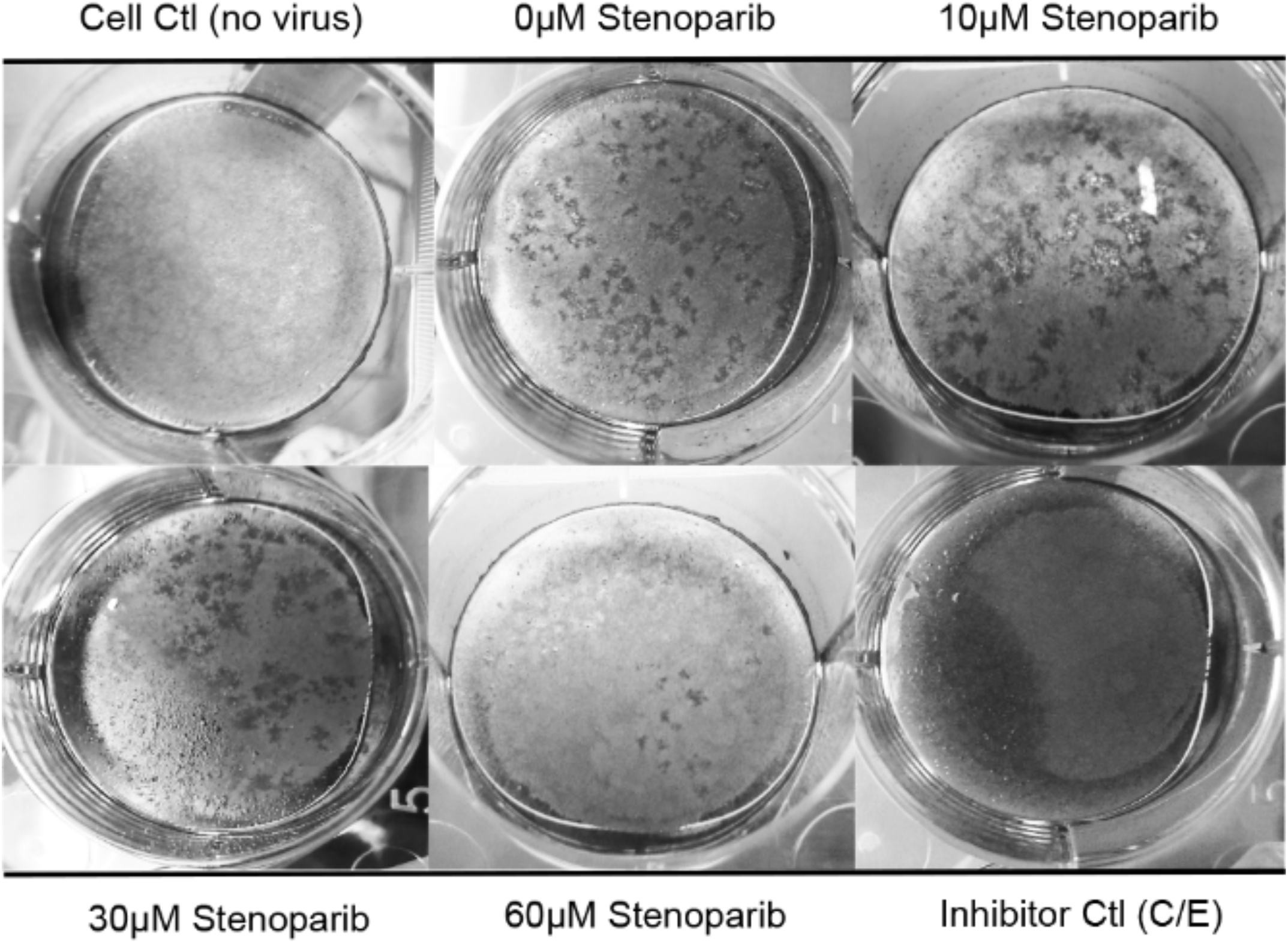
Stenoparib inhibits plaque formation in Calu-3 cells. Plaque assays were performed using Calu-3 cells infected with SARS-CoV-2 and treated with varying concentrations of stenoparib. Plaques are identified as empty regions or “dead zones” in the cellular monolayer and are expressed as plaque forming units (PFUs) per well. Plaques were manually counted and averaged among experimental replicates. This score is normalized as a percentage of untreated, but infected cells. In this representative image of the SARS-CoV-2/Calu-3 experiment, plaques are dark scars on the cellular monolayer. Controls were 1) uninfected cells (Cell Ctl), 2) untreated, but infected cells (0 μM), and 3) a camostat mesylate and E64d (C/E) inhibitor control (Inhibitor Ctl). Treatment with stenoparib at 10 μM and 30 μM led to a marked reduction in plaquing efficiency, whereas 60 μM resulted in near complete inhibition.

